# Donor Age and Oligosaccharide Structure Jointly Shape the Gut Microbiome Function

**DOI:** 10.64898/2026.06.07.730712

**Authors:** Ailing Zhang, Qing Wu, Hongye Qin, Janice Mayne, Zhibin Ning, Carolini Esmeriz da Rosa, Daniel Figeys

**Affiliations:** School of Pharmaceutical Sciences, Faculty of Medicine, University of Ottawa, Ottawa, ON, Canada; Department of Biochemistry, Microbiology and Immunology, Faculty of Medicine, University of Ottawa, Ottawa, ON, Canada; Laboratory of Nanobiotechnology and Applied Microbiology, Department of Food Science, Federal University of Rio Grande do Sul, Porto Alegre, Brazil; Quadram Institute Bioscience, Norwich Research Park, Norwich, Norfolk, United Kingdom; University of East Anglia, Norwich, Norfolk, United Kingdom

## Abstract

Non-digestible oligosaccharides are widely used as prebiotics, yet structurally related glycans can elicit distinct gut microbiome responses. Here, we combined controlled *ex vivo* fermentation, deep DIA metaproteomics, and targeted metabolomics to determine how oligosaccharide structure and donor age shape microbiome function. Stool microbiomes from 18 healthy donors across three age groups were cultured with seven structurally related oligosaccharides from two glycan families, fructo-oligosaccharides (FOS) and galactosyl-sucrose derivatives (GSD).

We found that oligosaccharide structure organized a functional response landscape rather than simply separating substrates into broad prebiotic classes. Structurally related glycans produced more similar response profiles overall, yet closely related FOS substrates remained functionally distinguishable, indicating that subtle structural differences were resolved by the microbiome as graded functional changes. These structure-responsive functions were further associated with producer-level reorganization relative to baseline, while targeted enzyme-level analyses indicated that substrate-specific CAZyme responses could also reflect altered functional investment within shared producer backgrounds. Despite these substrate-specific entry processes, network analysis revealed convergence onto shared downstream physiological states enriched for translation, amino-acid biosynthesis, secretion/export, and chemotaxis-related pathways. Across treatments, major short-chain fatty acids increased while mucin glycan degradation-associated markers decreased, suggesting coordinated shifts toward saccharolytic metabolism and reduced host-glycan foraging. Tryptophan-associated metabolism was also consistently linked to primary fructan processing, accompanied by higher extracellular tryptophan availability.

Donor age modified selected microbial functional axes and enzyme-metabolite coupling relationships rather than the overall direction of core fermentation outputs. In particular, oligosaccharides attenuated an *Methanobrevibacter smithii* (*M. smithii*) and M00567 methanogenesis-related signature in microbiomes from older adults and altered age-dependent relationships between butyrate-pathway enzymes and extracellular butyrate levels.

Together, these findings show that oligosaccharide structure determines how gut microbial communities organize carbohydrate processing and downstream functional states, while donor age reshapes the taxonomic and metabolic context of these responses. This work provides a mechanistic framework for structure-aware and age-aware precision prebiotic design.

## Introduction

Dietary non-digestible oligosaccharides are well-established modulators of the human gut microbiome. These short carbohydrate chains reach the colon intact, where they are selectively metabolized by resident microbes, stimulating the production of short-chain fatty acids (SCFAs) and shaping microbial community structure and metabolic output^1,2^. Fructo-oligosaccharides (FOS) and galactosyl-sucrose derivatives (abbreviated here as GSD in this study, including the raffinose family oligosaccharides and their β-linked isomers) represent two of the most prevalent classes of naturally occurring and commercially utilized prebiotics^3-5^. Although they share a sucrose core, they differ in glycosidic linkages, degree of polymerization, and overall branching, which together influence microbial accessibility, substrate preference, and the metabolic pathways engaged during fermentation^6,7^. Understanding how subtle structural differences among oligosaccharides determine microbial and metabolic responses remains an active area of research.

Microbial responses to prebiotics are not uniform across donors. Age is a major determinant of gut microbiome composition and function, and is linked to differences in community stability and recovery after perturbation^8-10^. Age-associated shifts in key taxa, including *Bifidobacterium, Bacteroides, Prevotella*, and butyrate-producing *Firmicutes*, can lead to changes in the ability of the microbiome to use specific prebiotics, the metabolic output of the microbiome and the ecological context in which a given substrate is fermented ^9,11,12^. As a result, the same oligosaccharide may engage different taxa and functional pathways across individuals, and age-related differences in baseline composition may also influence how functional changes relate to metabolite readouts^13-15^.

Few studies have systematically compared age-stratified responses to sets of structurally related oligosaccharides within a single experimental framework, and, to our knowledge, none have done so with the functional resolution provided by metaproteomics. Here, we used the RapidAIM^16^ ex vivo microbiome assay to test seven structurally distinct oligosaccharides, comprising four FOS-type and three GSD-type compounds, using fecal microbiota from 18 human donors spanning three age groups: young, mid, and old adults. Quantitative DIA-based metaproteomics was used to profile substrate-driven microbial functional changes in the microbiomes^17,18^ coupled with targeted metabolomics to profile specific changes in metabolites.

By integrating multi-age cohorts, structurally resolved oligosaccharides, metabolomic profiles, and deep metaproteomics within one unified system, this study provides a comprehensive view of how age and substrate jointly determine microbiome functional dynamics. The dataset offers mechanistic insights into carbohydrate utilization pathways, and individualized fermentation phenotypes, providing a foundation for precision prebiotic strategies tailored to donor-age and microbial ecology.

## Main Methods

### Sample collection and processing

Stool sample collections were approved by the University of Ottawa Office of Research Ethics and Integrity under protocols H-09-23-4985 and H-09-22-8466 (also the Bruyère Research Ethics Board (M16-19-039)) and by the Ottawa Health Science Network Research Ethics Board at the Ottawa Hospital (20160585-01 H). Eighteen gut microbiomes from healthy individuals aged 18–85 years were included in this study (Supplemental Table 1). Stool samples were self-collected using a validated anaerobic stabilization workflow and transported on cold packs. Upon receipt, samples were processed under anaerobic-stabilizing conditions, diluted to 20% (w/v), homogenized, clarified, filtered, and stored at −80°C until use.

### RapidAIM 3.0 *ex vivo* culture assay

Seven oligosaccharides were tested, including four fructo-oligosaccharides (FOS: KES, NYS, KP, and KH) and three galactosyl-sucrose/RFO-series oligosaccharides (GAL, RAF, and STA). PBS was used as the vehicle control. Based on the RapidAIM 2.0 platform^16^, individual microbiome samples were pre-incubated in culture medium for 6 h under anaerobic conditions, followed by treatment with oligosaccharides or PBS for an additional 12 h. Each treatment was performed in triplicate for each donor. After incubation, culture supernatants were collected for targeted metabolomics, and washed microbiome pellets were stored for metaproteomic analysis.

### Metaproteomic sample preparation and LC–MS/MS analysis

Microbiome pellets were processed using the SPEED^19^ protocol (Sample Preparation by Easy Extraction and Digestion) with minor modifications. Proteins were extracted, reduced, alkylated, and digested with trypsin. Peptides were desalted using reversed-phase tips on a robotic platform, dried, reconstituted, and analyzed by LC–MS/MS using a Vanquish Neo UHPLC system coupled to an Orbitrap Astral mass spectrometer. Data was acquired in data-independent acquisition mode.

### Database searching and functional annotation

DIA raw files from 486 samples were searched together using DIA-NN v2.3.0 with the InfinDIA module. The search database was the Unified Human Gastrointestinal Protein reference (UHGP v2.0.2) from MGnify^20^, containing 4,744 genomes. Trypsin was specified as the protease, allowing up to one missed cleavages. Carbamidomethylation of cysteine was set as a fixed modification. Quantification and cross-run normalization followed DIA-NN defaults. Output reports and expression matrices were filtered at 5% FDR at both precursor and protein-group levels. MetaX^21^ was used for downstream annotation and analysis.

### Targeted metabolomics

Post-RapidAIM culture supernatants were used for targeted metabolomics. Short-chain fatty acids (SCFAs), branched-chain fatty acids (BCFAs), and lactate were quantified using a 3-nitrophenylhydrazine derivatization workflow^22^ followed by UPLC–MS/MS. Amino acids were quantified using an aminoquinolyl-N-hydroxysuccinimidyl carbamate derivatization workflow^23^ on the same LC–MS/MS platform. Standard curves, blanks, and quality-control samples were included for quantitative assessment. Raw MS/MS spectra were processed in Skyline (v23.1)^24^ for peak integration and quantification.

## Data analysis

### Treatment response quantification

Treatment-associated functional responses were quantified as DESeq2^25^-derived log□ FC values relative to the matched PBS control. PBS was used as the reference condition for estimating treatment responses but was not included as an oligosaccharide in structure-based analyses. Unless otherwise specified, repeated measurements from the same donor were accounted for using linear mixed-effects models (LMMs) with donor as a random intercept. Multiple testing correction was performed using the Benjamini–Hochberg (BH) procedure.

### Primary CAZyme treatment-response modeling

For selected primary carbohydrate-active enzymes, including beta-fructosidase, beta-fructofuranosidase, alpha-galactosidase, and beta-galactosidase, treatment responses were modeled using DESeq2-derived log□FC values relative to PBS. Within each oligosaccharide series, treatments were encoded as ordered levels and modeled using LMMs with a spline term for treatment order and donor as a random intercept. Overall treatment-associated trends were assessed by likelihood ratio tests comparing the full model with a null model lacking the spline term. Treatment-specific estimated marginal means and 95% confidence intervals were used to summarize response patterns, and post hoc treatment contrasts were adjusted for multiple comparisons.

### Glycan structural distance and functional response distance analysis

To test whether oligosaccharide structure was associated with functional response divergence, pairwise glycan structural distances were compared with donor-level functional response distances. Structural distances were calculated using Gower distance based on curated glycan features, including degree of polymerization, monosaccharide composition, and galactosyl linkage. For each donor, DESeq2-derived log□FC responses relative to PBS were standardized across oligosaccharides, and pairwise Euclidean distances between glycan response profiles were calculated using features with complete response estimates. Associations between structural and functional distances were tested using the model: Functional distance ∼ Structural distance + (1 | donor). Statistical support was further evaluated using glycan-label permutation tests, donor-level Mantel tests, and sensitivity analyses using alternative distance definitions and within-family glycan subsets.

### FOS degree of polymerization response modeling

FOS chain-length-associated responses were tested using DESeq2-derived log□FC values relative to PBS. KES, NYS, KP, and KH were encoded by degree of polymerization (DP = 3–6), and KEGG module- and KO-level responses were modeled separately using LMMs: log□FC ∼ DP + (1 | donor). The total response gradient across the FOS series was defined as the model-estimated difference between DP6 and DP3, with confidence intervals calculated on the same scale. P values were adjusted across features using the BH procedure. Module-level results were used to summarize broad functional gradients, while significant KO-level positive and negative response sets were used for producer-guild analysis.

### Producer-guild analysis of FOS DP-responsive KOs

To determine whether FOS DP-responsive functions were associated with distinct producer organization, significant KO-level results from the FOS DP response model were separated into positive and negative response-gradient sets. For each donor and FOS treatment, producer profiles were constructed by aggregating the abundances of species contributing to KOs within each response set. Producer composition shift was quantified as the Bray–Curtis distance between each treatment-associated producer profile and the matched PBS producer profile from the same donor. To account for differences in KO set size, size-matched resampling was used to compare the negative response set against a null distribution generated from the larger positive response set. Empirical P values were calculated from the resampling distribution and adjusted across FOS treatments using the BH procedure.

### Co-expression module and pathway enrichment analysis

KO co-expression modules were constructed using weighted gene co-expression network analysis (WGCNA)^26^ based on log-transformed KO abundance profiles. Sparse KOs were filtered before network construction, and signed networks were built using robust correlation. Modules were identified by hierarchical clustering of the topological overlap matrix and dynamic tree cutting, followed by merging of closely related modules based on eigengene similarity. For each target primary CAZyme, the module with the highest module membership was selected for downstream interpretation. KEGG pathway gene set enrichment analysis was then performed using signed module membership values as the ranking statistic. Enrichment significance was assessed using BH correction, and normalized enrichment scores were used to summarize enrichment direction and magnitude.

### Metabolite and KO–SCFA association analysis

Metabolite responses were analyzed separately for each experimental round using LMMs. Measurements below the detection limit were set to zero, background-corrected using matched medium-only controls, transformed using the inverse hyperbolic sine transformation, and averaged across technical replicates for each donor. Age-group effects were assessed using the model: y ∼ Treatment × AgeGroup + (1 | donor). A complementary continuous-age analysis was performed using mean-centered age: y ∼ Treatment × Age_c + (1 | donor). For each metabolite, age, treatment, and interaction effects were assessed by likelihood ratio tests on nested models, with BH correction across metabolites within each round.

For selected KO–SCFA pairs, associations were tested using log□-transformed KO and SCFA abundances. Models were fitted separately within the FOS and GSD rounds: SCFA ∼ KO × AgeGroup × Treatment + (1 | donor). Age- and treatment-specific KO–SCFA slopes were estimated using marginal trends, and slope-level P values and pairwise slope contrasts were adjusted using the BH procedure.

### Age-associated taxa and M00567 methanogenesis-related analysis

Age-associated differences in PBS-only taxon intensities were analyzed using LMMs after averaging technical replicates. Taxa with sufficient detection across donors were modeled as: log□ (Abundance + c) ∼ AgeGroup + PBS_Rep + Sex + (1 | donor). where c was defined as half of the smallest positive abundance observed for each taxon. Overall age effects and pairwise age-group contrasts were corrected using the BH procedure. For methanogenesis-related functional analysis, KEGG module M00567 was analyzed separately within each oligosaccharide series. PBS age-group differences were tested using Tukey-adjusted pairwise contrasts, and treatment-associated changes were evaluated within old donors using LMMs with donor as a random intercept and Dunnett-adjusted contrasts against PBS.

Detailed experimental and computational methods are provided in the Supplemental methods.

## Results

### Glycan structural distance predicts functional response divergence

Briefly, RapidAIM assays were performed across 18 donor microbiomes, seven oligosaccharide treatments, and matched PBS controls, yielding 486 culture samples with three technical replicates per donor-condition combination and identifying 1,788 taxa and 181,214 peptides across all samples. This design enabled us to ask whether structural differences among oligosaccharides were systematically reflected in microbiome functional responses. We therefore compared pairwise glycan structural distances with donor-level KEGG module response distances. The structural distance matrix separated the four FOS substrates from the three GSD substrates while also preserving within-family gradients, particularly across the FOS chain-length series (Fig. 2a). Consistent with this structure, pairwise functional response distances were lowest among closely related FOS substrates, intermediate among GSD substrates, and highest between FOS and GSD glycans (Fig. 2b).

Across donor-level pairwise comparisons, functional response distance increased strongly with glycan structural distance (LMM slope = 0.899; Fig. 2c). This association remained significant after glycan-label permutation testing (permutation *p = 0*.*0035*), indicating that the observed relationship depended on the actual glycan structural labels rather than the pairwise data structure alone. Subject-level Mantel analyses further showed that this structure–function concordance was consistent across individuals, with mostly high positive correlations and a median Mantel r of 0.761 (Fig. 2d; Supplemental Table 2). These results indicate that structurally related oligosaccharides induced more similar KEGG module response profiles, whereas structurally divergent glycans induced more distinct functional responses.

Having established this global relationship, we next asked whether a specific structural feature within the FOS series, degree of polymerization, was associated with module-level response gradients. Modeling responses from DP3 to DP6 identified 68 significant KEGG modules, including 53 with positive DP-associated effects and 15 with negative effects (*q < 0*.*05*, Supplemental Table 3). Positive DP-associated gradients were more frequent than negative gradients and were enriched among biosynthetic and central metabolic modules, whereas negative gradients involved a smaller and more heterogeneous set of metabolic pathways (Fig. 2e). Most effects were modest in magnitude, suggesting that closely related FOS substrates were not functionally interchangeable, but instead produced broad and graded remodeling of module responses. Together, these analyses show that oligosaccharide structure organized the overall functional response landscape across oligosaccharides, while closely related FOS substrates remained functionally distinguishable through predominantly small but significant graded module-level responses.

### Fructan-cleaving CAZymes correlate with tryptophan biosynthesis despite limited overlap in producer taxa

We explored whether there are correlations between the abundance of CAZymes involved in cleaving glycosidic bonds and other enzymes. In particular, Fructan beta-fructosidase (short as fructosidase below) and beta-fructofuranosidase (short as fructofuranosidase below), two enzymes involved in cleaving glycosidic bonds releasing terminal fructose units from fructosyl-containing oligosaccharides, but differ primarily in substrate specificity. As shown in the Venn diagram (Fig. 3a, d), tryptophan synthase was consistently and positively correlated with all four groups (*bicor* > 0.4).

To assess whether the bacteria encoding the fructan-releasing CAZymes overlapped with those producing tryptophan synthase, we examined the taxonomic origins of two key fructan-cleaving enzymes, fructosidase and fructofuranosidase. Across both FOS and GSD treatments, fructosidase was dominated by members of Bacteroidaceae and Tannerellaceae, spanning four Bacteroidota genera (*Bacteroides, Phocaeicola, Parabacteroides*, and *Prevotella*), with an additional contribution from *Anaerostipes* (Lachnospiraceae). In contrast, fructofuranosidase was primarily assigned to Lachnospiraceae, Ruminococcaceae, Bifidobacteriaceae, Coriobacteriaceae, and Erysipelatoclostridiaceae, and its producers were largely distinct from those contributing fructosidase. At the species level, *Bacteroides thetaiotaomicron* and *Anaerostipes hadrus_A* encoded both enzymes, whereas *Anaerostipes sp900066705* was consistently assigned only fructosidase (Fig. 3b). Notably, FOS recruited a broader β-fructofuranosidase producer repertoire than GSD, extending to additional genera and species beyond the shared core set.

We then profiled the producers of tryptophan synthase. Across both FOS and GSD treatments, tryptophan synthase was assigned to seven families, comprising 13 genera and 14 species. Four of these families overlapped with the CAZyme-producing families (Bacteroidaceae, Tannerellaceae, Lachnospiraceae, and Bifidobacteriaceae), whereas Erysipelotrichaceae, Enterobacteriaceae, and Anaerotignaceae were unique to tryptophan synthase (Fig. 3c). Despite family-level overlap, only three species (*Prevotella copri_A, Phocaeicola dorei*, and *Parabacteroides distasonis*) encoded both tryptophan synthase and fructosidase. In contrast, none of the fructofuranosidase producers were assigned as tryptophan synthase producers under either treatment (Fig. 3d).

We further quantify the consistency of tryptophan biosynthesis response across donors, through per-subject DESeq2 results for tryptophan synthase across all oligosaccharide treatments relative to PBS. Tryptophan synthase showed a robust and donor-consistent up-regulation for all treatments except KES (94% of individuals exhibited a significant increase (*Padj* < 0.05)), whereas RAF induced significant up-regulation in all 18 donors (Fig. 3e). In contrast, tryptophanase was down-regulated in nearly all donors across treatments (Fig. 3e). Consistent with these proteomic signatures, extracellular tryptophan concentrations were higher globally across oligosaccharide treatments than under PBS (Fig.5c).

### Oligosaccharide structure shapes primary CAZyme programs and producer-guild organization

We first examined whether oligosaccharide chain length influenced the abundance of the main fructan-cleaving CAZymes in the FOS set (Fig. 4a; Fig 1). Using DESeq2-derived log□ fold changes relative to PBS as the response in linear mixed-effects models, both fructosidase and fructofuranosidase showed significant treatment effects (LRT *p* ≈ 1.19 × 10^−1^□ and 7.16 × 10^−^ □, respectively). Across the FOS series, both enzymes showed progressively stronger induction with increasing chain length, although the response was more pronounced for fructosidase (Fig 4a). Tukey-adjusted pairwise contrasts supported this overall pattern: fructosidase was less induced by KES than by NYS, KP, or KH, and was also lower under NYS than under KH, whereas NYS and KP, and KP and KH, did not differ significantly. For fructofuranosidase, only the contrasts between KES and KP and between KES and KH were significant (Fig 4c). Under equal-mass addition, the total supplied fructose residues increased only modestly across the KES-to-KH series; comparison with residue-based expectations (Supplemental Table 5) showed that the observed separation was larger than expected from fructose-residue input alone, particularly for fructosidase, while fructofuranosidase showed a weaker but directionally similar departure that was most evident in contrasts between KES and the longer-chain treatments (Supplemental Fig. 1). Together, these results indicate a coherent structure-dependent response across the FOS substrates.

**Figure 1.**
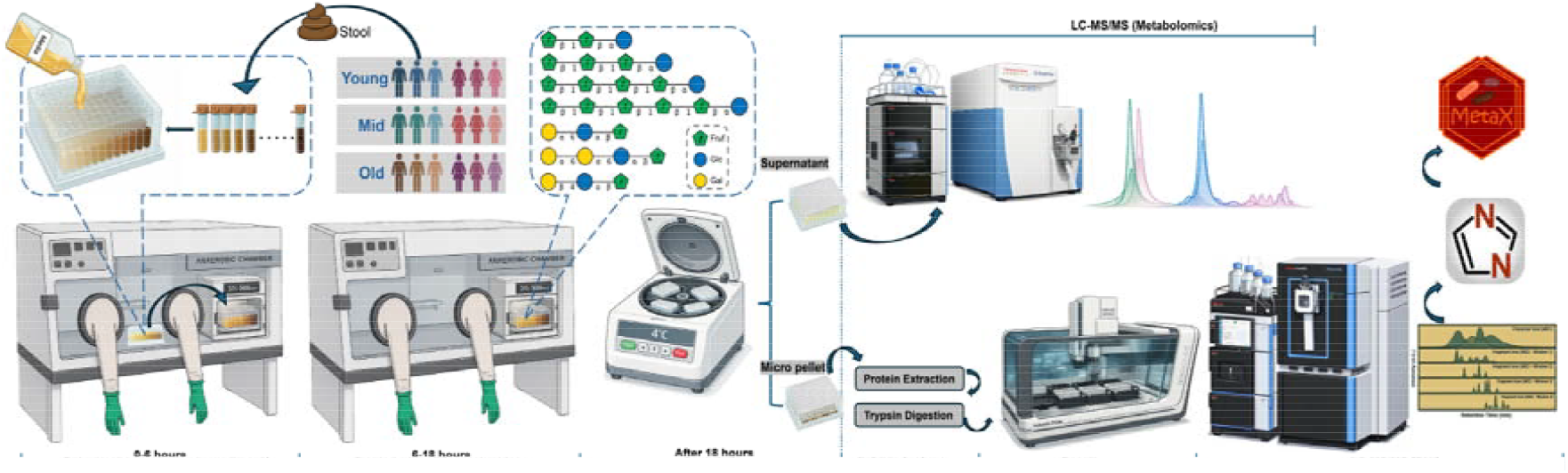
Pipeline of workflow. Biologic description of oligosaccharides from PubChem^27^.

**Figure 2.**
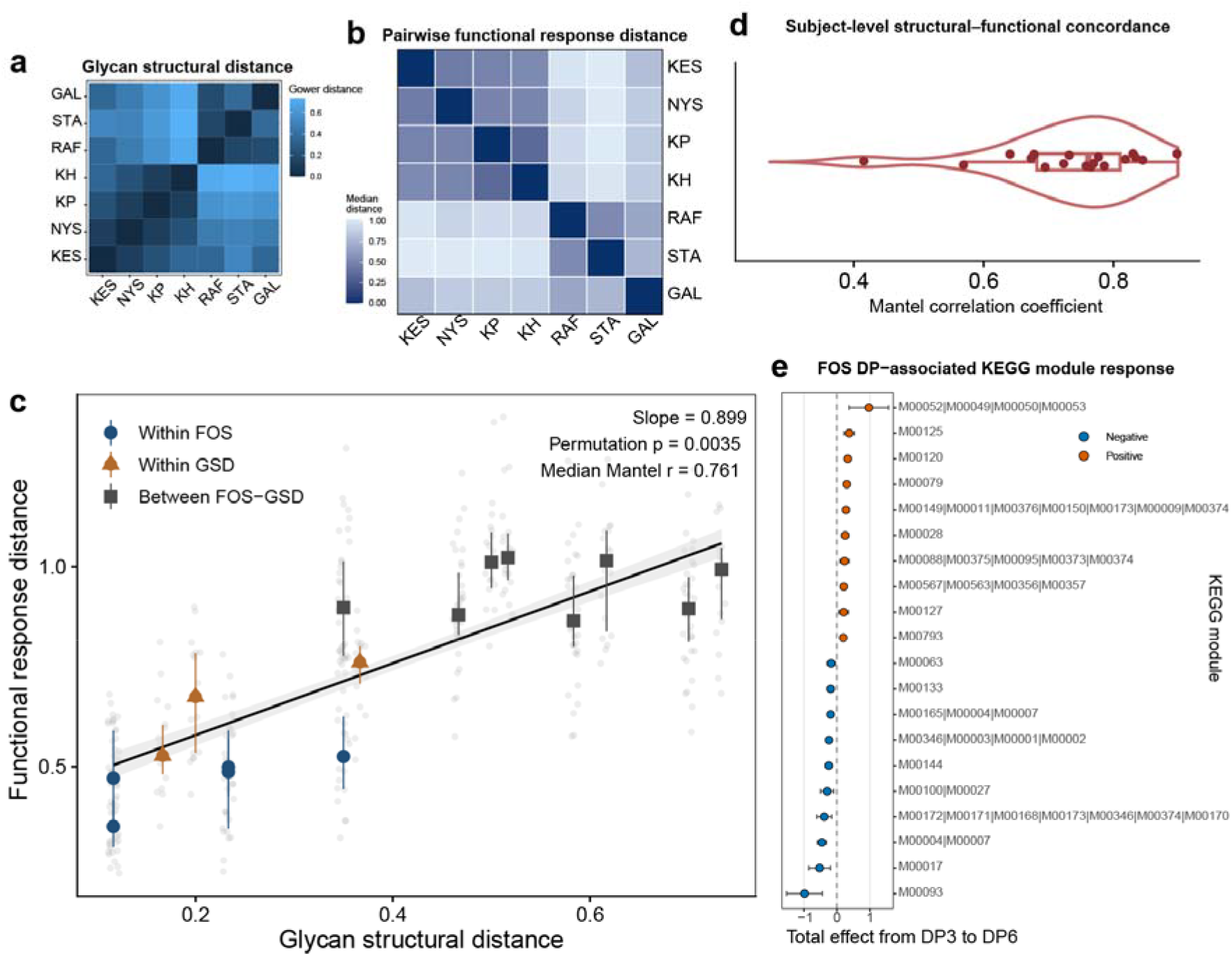
Glycan structural distance predicts functional response divergence and FOS DP-associated module gradients. (a) Pairwise glycan structural distances among the seven oligosaccharides, calculated using Gower distance from curated structural features including degree of polymerization, monosaccharide composition, fructan linkage, and galactosyl linkage. PBS was used only as the reference for response calculation and was not included as a glycan. (b) Pairwise functional response distances calculated from standardized KEGG module log_2_FC responses using Euclidean distance and summarized as the median across donors. (c) Relationship between glycan structural distance and donor-level functional response distance. Grey points represent donor-level pairwise glycan comparisons. Colored points summarize glycan-pair classes: within FOS, within GSD, and between FOS and GSD. The fitted line shows the mixed-effect model estimate with 95% confidence interval. Slope, glycan-label permutation p value, and median subject-level Mantel r are shown. (d) Subject-level Mantel correlations between glycan structural distance and KEGG module response distance. Each point represents one donor. (e) FOS degree-of-polymerization-associated KEGG module response gradients from DP3 to DP6. Points show total estimated effects, and horizontal bars indicate 95% confidence intervals. Positive and negative DP-associated modules are shown in orange and blue, respectively. Only module IDs are shown.

**Figure 3.**
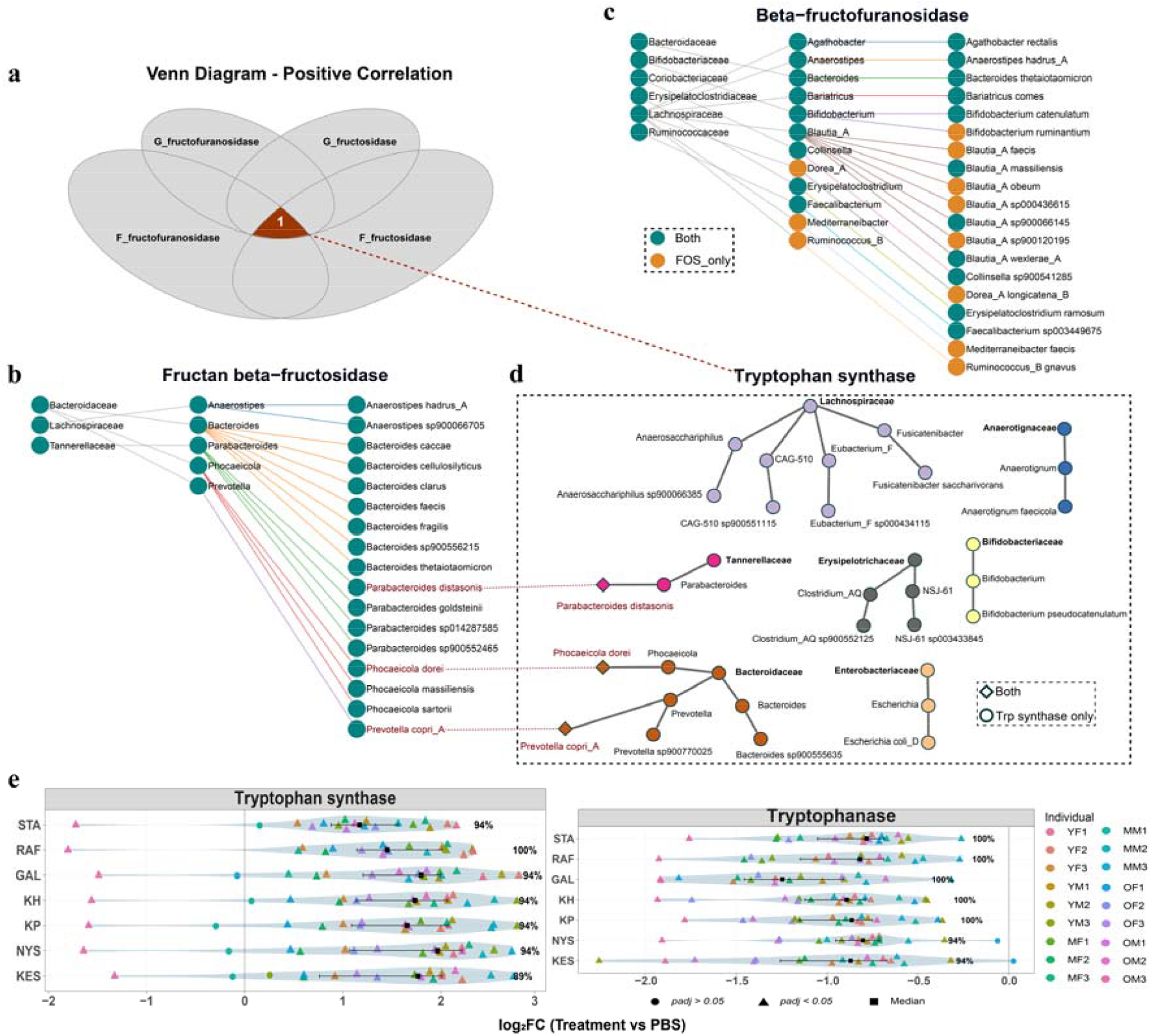
Co-expression context and community sources of fructan-degrading CAZymes and tryptophan metabolism across oligosaccharide treatments. (a) For the two first-cleaving CAZymes: fructan β-fructosidase and β-fructofuranosidase EC-level co-expression was computed separately within the FOS and GSD treatment series using bicor. For each target enzyme and each series, we collected enzymes showing positive correlation with *bicor > 0*.*4* and BH-adjusted *p < 0*.*05*, and summarized the overlap among the four sets (FOS vs GSD× two target enzymes) using a Venn diagram. (b–c) Taxonomic attribution of proteins annotated as fructan β-fructosidase (b) and β-fructofuranosidase (c). Node color indicates whether a taxon was detected under both FOS and GSD series or FOS-only (as labeled). (d) Taxonomic attribution of tryptophan synthase producers. Symbols indicate taxa encoding both tryptophan synthase and fructan-hydrolyzing CAZymes versus taxa encoding tryptophan synthase only (as labeled in the panel). (e) Per-individual response of tryptophan synthase (top) and tryptophanase (bottom) under each treatment, shown as log_2_FC (Treatment vs PBS) for each subject (colored points; subject IDs listed at left). Black squares denote the median across individuals within each treatment. Point shape indicates significance. Percentages at the right indicate the proportion of individuals showing a response in the dominant direction for that feature under the corresponding treatment.

**Figure 4.**
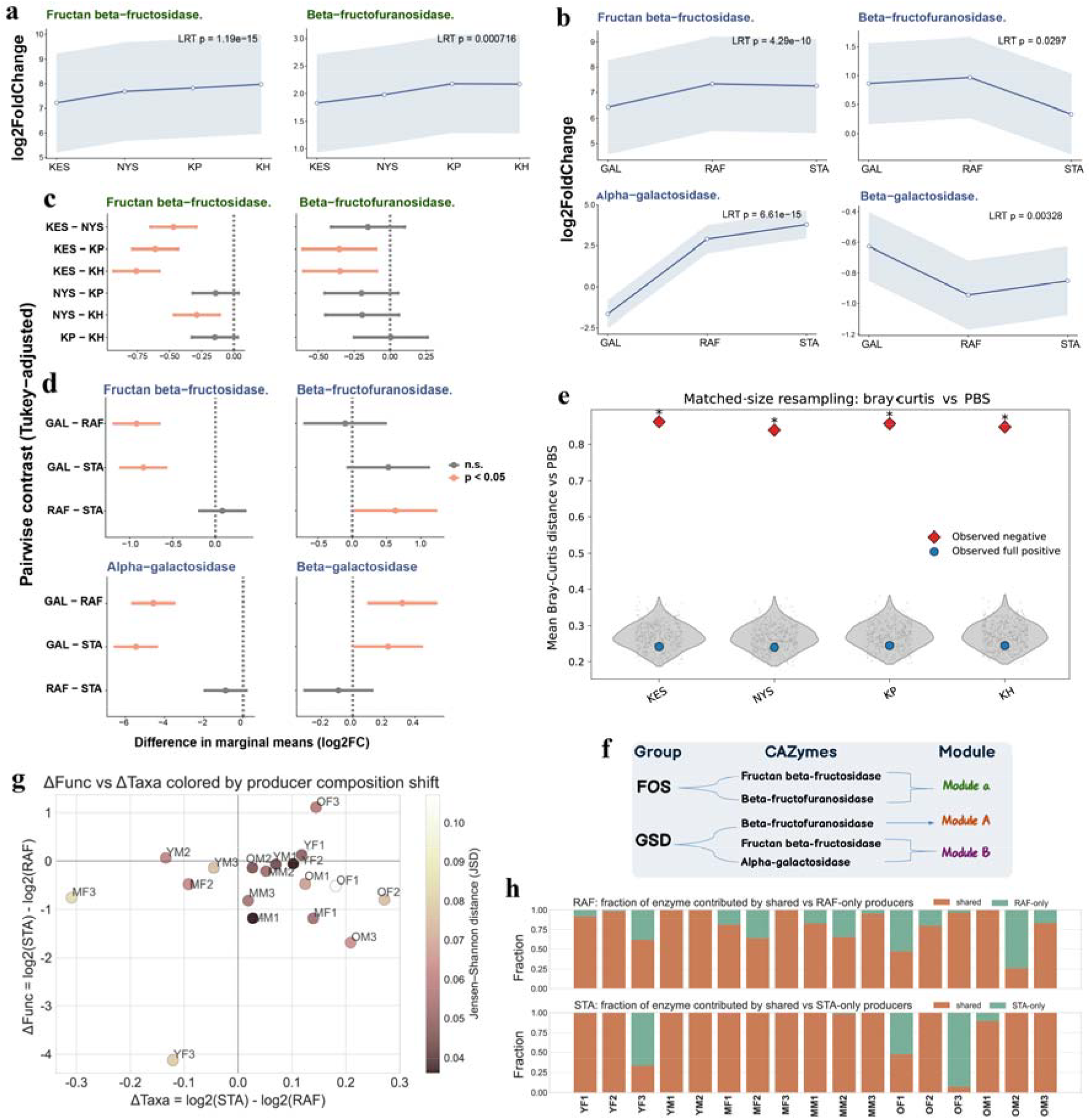
Primary CAZyme responses and producer-level organization across oligosaccharide treatments. (a-b) DESeq2-derived log□ fold-change values relative to PBS were modeled using linear mixed-effects models, and treatment effects were evaluated using likelihood ratio tests. Lines represent estimated marginal means, and shaded areas show 95% confidence intervals. (c–d). Tukey-adjusted pairwise contrasts of CAZyme across FOS and GSD treatments. Differences in estimated marginal means (log□ fold-change relative to PBS). Points indicate contrast estimates, and horizontal bars represent 95% confidence intervals; colored points denote adjusted *p < 0*.*05*. a/c: FOS treatments, b/d: GSD treatments. (e) Size-matched resampling analysis of producer compositional shift for FOS DP-associated KO response sets. Bray–Curtis distance was calculated between each treatment-associated producer profile and the matched PBS profile. Grey violins and points show null distributions generated by random positive KO subsets matched to the size of the negative KO set. Red diamonds indicate the observed negative KO set, and blue circles indicate the observed full positive KO set. Asterisks indicate significant deviation of the observed negative set from the size-matched positive null distribution. (f) Schematic diagram of the modules showing the highest kME for the target CAZymes. (g) Each point represents one subject. The x-axis shows the change in total producer abundance between STA and RAF. The y-axis shows the enzyme-level change derived from DESeq2. Points are colored by the Jensen–Shannon distance (JSD) between the normalized producer contribution profiles under RAF and STA, where larger values indicate greater reweighting of producer contributions. (h) Producers were partitioned into shared (active in both RAF and STA) and treatment-only sets using species-level contributions. Total contribution was decomposed accordingly, and shared fractions were computed for each treatment.

Because the FOS series showed a coherent degree-of-polymerization-associated response, we next asked whether the KO-level FOS DP-associated response sets identified above were linked to distinct producer-guild organization. We used the positive and negative DP-associated KO sets from the structure–response analysis as inputs for producer-level attribution and compared their producer compositional shifts from matched PBS (Supplemental Table 6). To account for the larger size of the positive KO set, we performed size-matched resampling by repeatedly sampling positive KO subsets matched to the size of the negative KO set. Across all FOS treatments, the observed negative KO set showed a much larger Bray–Curtis shift from matched PBS than expected from the size-matched positive null distributions, whereas the full positive KO set remained close to the size-matched positive range (Fig. 4e). These results indicate that the negative FOS DP-associated KO responses were linked to a distinct producer compositional shift that was not explained by KO set size imbalance.

In the GSD set, fructofuranosidase displayed a non-monotonic response, peaking under RAF and declining under STA; accordingly, RAF versus STA was the only significant pairwise contrast, whereas GAL did not differ significantly from either substrate (Fig. 4b). Comparison with residue-based expectations under equal-mass addition further showed that the RAF–STA separation for fructofuranosidase was larger than expected from fructose-residue input alone (Supplemental Fig. 2; Supplemental Table 7). By contrast, alpha-galactosidase increased strongly across substrates containing α-linked galactose units (LRT p ≈ 6.61 × 10^−1^□) (Fig. 4b), with GAL showing lower abundance than both RAF and STA (Fig. 4d). The RAF–STA difference for alpha-galactosidase was broadly consistent with the higher galactose-residue supply under STA (Supplemental Fig. 2; Supplemental Table 7). Beta-galactosidase showed the opposite trend and was consistently downregulated across the GSD treatments (LRT p ≈3.28 × 10^−3^), including under GAL, with the strongest decreases observed under RAF and STA (Fig. 4b, d). Together, these results indicate that GSD substrates elicit more differentiated first-cleavage CAZyme responses than the largely coherent trend observed across the FOS series.

To place these first-cleaving CAZymes into broader functional contexts, we next performed WGCNA separately in the FOS and GSD datasets using KO-level intensities across samples. In the FOS dataset, fructosidase and fructofuranosidase showed their strongest associations with the same module context, consistent with their coordinated response patterns across the FOS series (Fig. 4f, Supplemental Table 8). In the GSD dataset, alpha-galactosidase and fructosidase likewise showed their strongest associations with the same module context, whereas fructofuranosidase was most strongly associated with a different one (Fig. 4f, Supplemental Table 9). Although the lower fructofuranosidase signal under STA than RAF may partly reflect the lower β-fructose content in STA when compounds were added at equal total amounts, comparison with residue-based expectations showed that the RAF–STA difference exceeded that predicted from fructose-residue input alone. Together with the distinct module association of fructofuranosidase in the GSD network, these observations motivated us to ask whether the STA-associated reduction could be explained by changes in the abundance or composition of linked producer species.

We therefore asked whether this deviation could be accounted for by changes in the summed abundance or composition of linked producer species. For each donor, we compared the STA–RAF change in fructofuranosidase intensity (ΔFunc) with the corresponding change in the summed abundance of linked producer species (ΔTaxa). Although fructofuranosidase was lower under STA than RAF in 15 of 18 donors, the summed abundance of its linked producers was higher under STA in 13 of 18 donors. Consistently, the residual term (ΔR = ΔFunc − ΔTaxa) was negative in 15 of 18 donors. These results indicated that the lower fructofuranosidase signal under STA was not accompanied by a corresponding decrease in the summed abundance of linked producers (Fig. 4g; Supplemental Fig. 3).

We therefore next asked whether this pattern reflected a shift in producer composition between treatments rather than a reduction in total producer abundance. For each donor, we partitioned fructofuranosidase abundance into contributions from producers shared between RAF and STA and from treatment-specific producers. In most donors, the fructofuranosidase signal under STA was dominated by shared producers, and only two donors showed substantial contributions from STA-specific species (Fig. 4h). Together, these findings suggest that the lower fructofuranosidase signal under STA was not driven by broad replacement of enzyme-associated producers, but instead reflected abundance-independent functional shifts within a largely shared producer background.

### Convergent module enrichment supports an anabolic shift, while mucin glycan degradation-associated markers show structure-dependent suppression

To interpret the functional themes most strongly associated with the target CAZymes, we performed KEGG pathway GSEA separately for the FOS and GSD modules showing the highest kME association with the respective target CAZymes, using WGCNA kME as the ranking metric. Although the two modules were derived from independent analyses and were associated with distinct oligosaccharide classes, they displayed broadly similar pathway enrichment profiles and concordant NES directions (Fig. 5a–b). In both modules, positively enriched pathways were dominated by translation- and biosynthesis-related functions, including Ribosome, Aminoacyl-tRNA biosynthesis, Biosynthesis of amino acids, and Valine, leucine and isoleucine biosynthesis, together with related central metabolic pathways such as 2-oxocarboxylic acid metabolism. Pathways related to protein trafficking and bacterial interaction, including Protein export, Bacterial secretion system, and Bacterial chemotaxis, were also positively enriched in both modules. By contrast, negatively enriched pathways in both modules included functions related to lipid metabolism, amino-acid catabolism, and carbohydrate interconversions, such as Fatty acid degradation, Glycerolipid metabolism, Lysine degradation, Tyrosine metabolism, and Pentose and glucuronate interconversions (Fig. 5b). Taken together, these shared enrichment trends are consistent with a common carbohydrate-responsive, growth-associated functional program, despite the distinct upstream substrate structures and their associated core CAZymes.

**Figure 5.**
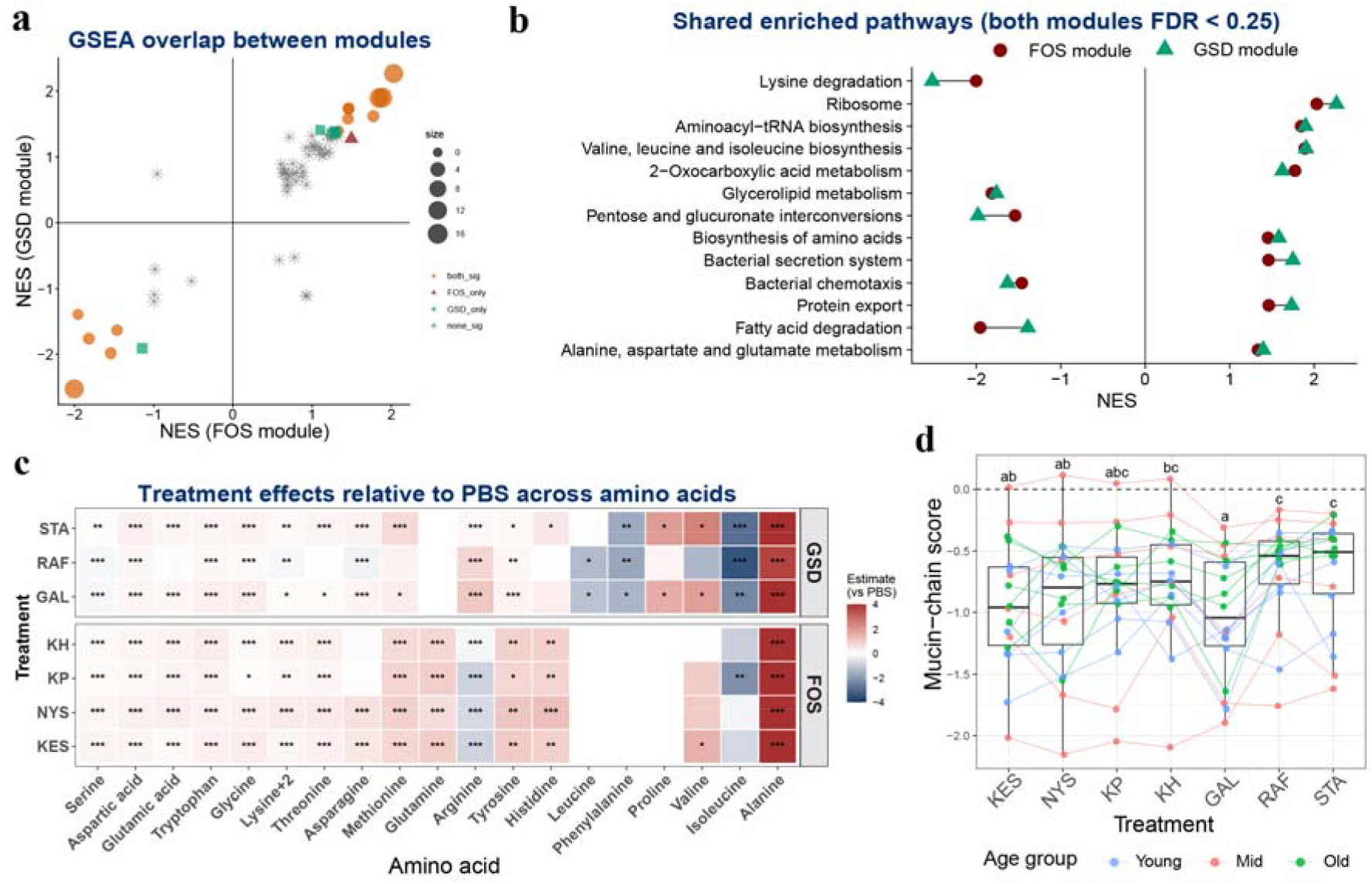
Functional interpretation of CAZyme-associated modules and downstream metabolic profiles. (a–b) GSEA was performed to functionally interpret the WGCNA modules showing the highest kME with the target CAZymes (FOS: blue module; GSD: green module), using KO lists ranked by kME. (a) NES concordance scatter plot for pathways shared between the two module-level GSEA results; point size scales with -log_10_(max (FDR_FOS_, FDR_GSD_)) and reference lines indicate NES = 0. (b) Dumbbell plot showing term-wise paired NES values for shared significant pathways/terms (*FDR < 0*.*25* in both modules), ordered by max (FDR_FOS_, FDR_GSD_); the vertical line marks NES = 0. (c) Tile color indicates the estimated treatment effect relative to PBS from emmeans contrasts based on LMM fitted to mean asinh-transformed net amino acid levels. Asterisks indicate BH-adjusted significance levels: *\*P < 0*.*05, **P < 0*.*01, ***P < 0*.*001*. For visualization, the color scale was truncated at ±4. (d) Subject-level mucin-chain scores across treatments. Each point represents one subject (colored by age group), and grey lines connect repeated measures from the same subject across treatments. Boxplots summaries the distribution across subjects for each treatment. The dashed horizontal line indicates no change relative to PBS (log_2_FC = 0). Different letters indicate significant differences between treatments and treatments sharing at least one letter are not significantly different (α *= 0*.*05*).

Because amino acid biosynthesis and metabolism pathways were enriched in modules associated with the primary CAZymes, we quantified 19 amino acids in the culture supernatant and summarized their treatment-versus-PBS effects in a heatmap (Fig. 5c). In both oligosaccharide sets, most amino acids showed positive effects relative to PBS. In the GSD set, several amino acids, including lysine, serine, and asparagine, displayed more treatment-dependent patterns. By contrast, the FOS set showed more similar response profiles across treatments, with arginine showing an opposite pattern to that observed in the GSD set. Notably, although many amino acids were higher under oligosaccharide treatments than under PBS, their levels generally remained below the amino acid background present in the starting medium, indicating reduced net consumption and oligosaccharide treatments may shift community metabolism away from amino acid catabolism (Supplemental Fig. 4). No significant age-associated changes were detected.

To characterize treatment-dependent shifts in mucin glycan degradation-associated markers, we analysed a panel spanning desulfation and terminal de-capping (arylsulfatase type I, GH33, GH29, and GH95), endo release (GH16_3), exo trimming (GH2, GH35, GH20, and GH84), core dismantling (GH101), and side-chain processing, including ABO-related branches (GH36, GH109, and GH110) and type-1 chain processing (GH136)^28^. A subject-resolved heatmap of log_2_FC (treatment vs PBS) showed broadly negative shifts across multiple mucin-associated functions, although the degree of coordination varied across markers (Supplemental Fig. 5a). Consistent with this pattern, the fraction of subjects with negative log_2_FC values was highest for GH20, GH136, GH109, and GH33, and was weak or inconsistent for GH84 (Supplemental Fig. 5b). Overall, these results indicate a coordinated reduction across several key mucin-foraging functions, but not a uniform shift across the entire marker panel.

We next summarized these coordinated changes at the subject level using the mucin-chain score. Scores were negative under all treatments, indicating an overall downward shift relative to PBS. CLD and Tukey-adjusted pairwise comparisons showed the strongest downshifts under GAL, KES, and NYS, the weakest under RAF and STA. Specifically, GAL was significantly lower than KH, RAF, and STA, whereas KES and NYS were significantly lower than RAF and STA. All other pairwise differences were not significant (Fig. 5d).

### Oligosaccharide treatment reduces an age-associated *M. smithii* and M00567 signature

Overall, taxonomic composition showed strong donor-specificity, where samples from the same donor generally retained similar compositional structures despite different treatments. The results are reproducible across independent experimental analyses and across distinct oligosaccharide treatments (Supplemental Fig. 6). At the phylum level, Bacteroidota and Firmicutes_A represented the dominant fractions in most donors. Proteobacteria and Actinobacteriota also contributed substantial proportions in some individuals. At the genus level, *Anaerostipes, Bacteroides, Blautia_A, Faecalibacterium, Parabacteroides*, and *Phocaeicola* were among the most abundant genera across donors. However, the relative abundances of these dominant genera varied across individuals, and treatment-associated shifts were not uniform among donors.

**Figure 6.**
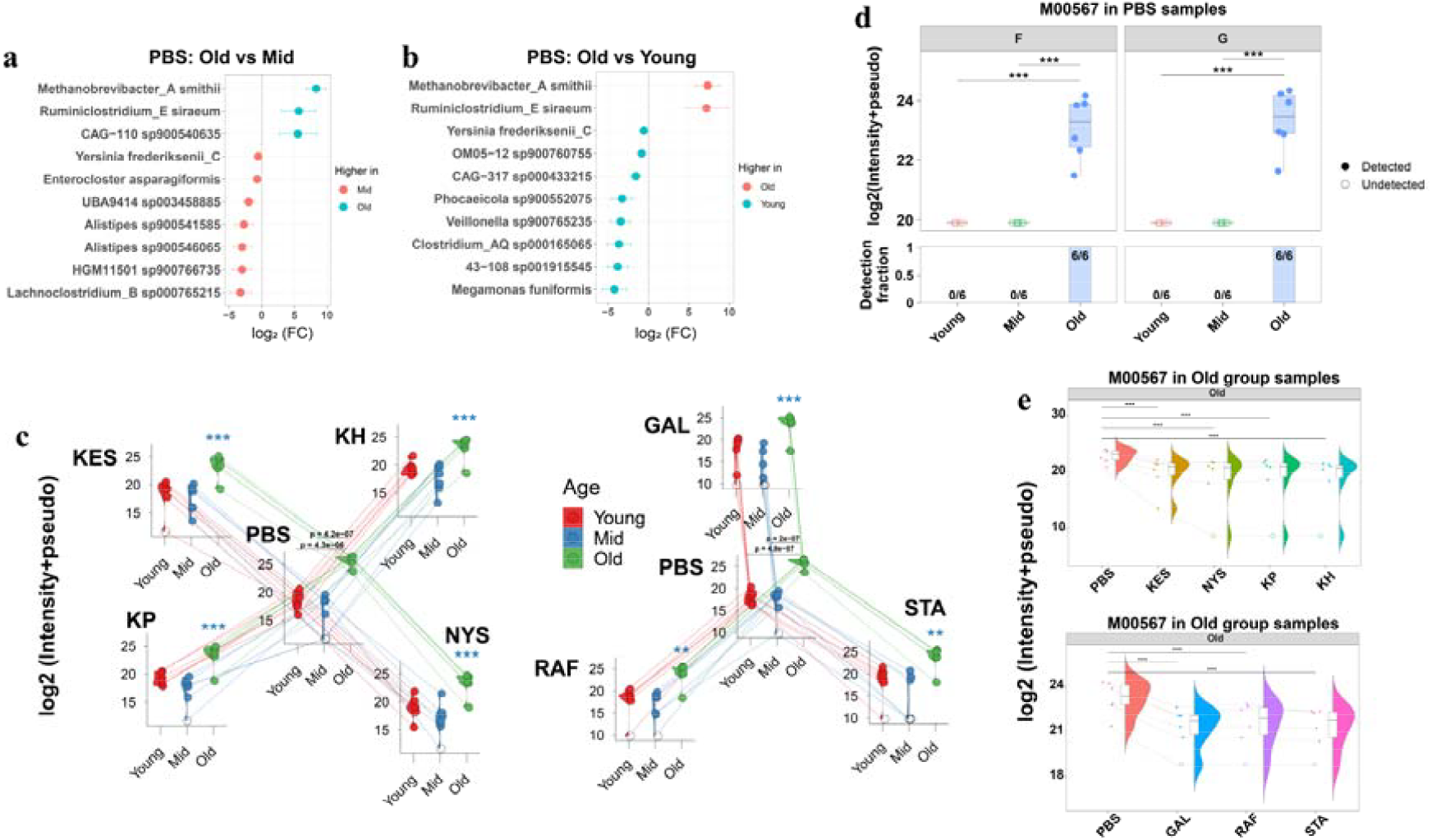
Age-associated *M. smithii* enrichment and oligosaccharide-associated reduction of M00567 in older microbiomes. (a–b) Age-associated taxa identified by LMM and validated by LOSO. Shown are the top 10 species for (a) old vs mid and (b) Old vs young age groups, ranked by FDR. Points indicate log_2_FC, with horizontal bars showing 95% confidence intervals. (c) Log□ intensities of *M. smithii* across PBS and oligosaccharide treatments. Each point represents one donor-level sample, and lines connect paired PBS and treatment values from the same donor within each age group. (d) PBS abundance of KEGG module M00567, annotated as CO□-to-methane methanogenesis, across age groups in the FOS and GSD datasets. Solid points indicate detected values, and open circles indicate undetected values plotted at the pseudocount floor. The lower bar plot shows the detection fraction, with labels indicating detected samples over total samples in each age group. (e) Treatment-associated changes in M00567 abundance within old microbiomes for FOS and GSD treatments relative to PBS. For panels c and e, treatment-associated changes were tested within each age group using LMMs with donor as a random effect, followed by Dunnett-adjusted contrasts against PBS. For the PBS age-group comparisons in panels c and d, one-way models with emmeans-based Tukey-adjusted pairwise contrasts were used. Asterisks indicate significance (*\*\*P < 0*.*01*; *\*\*\*P < 0*.*001*).

To explore potential taxonomic differences between different age groups, we next examined taxonomic intensity under the PBS condition, and species-level comparisons identified taxa that differed between age groups by LMM. Across age-group, there was limited overlap in species; however, two species (*Methanobrevibacter smithii* (GTDB^29^: *Methanobrevibacter_A smithii* short as *M. smithii*) and *Ruminiclostridium_E siraeum*) were consistently enriched in the old group, and in particular *M. smithii* with a strong statistical significance (Fig. 6a, b, c; Supplemental Fig. 7). Interestingly, the abundance of *M. smithii* was consistently reduced following the treatment of the microbiomes with the prebiotics in the older age groups which was not the case in the mid or young groups (Fig. 6c).

**Figure 7.**
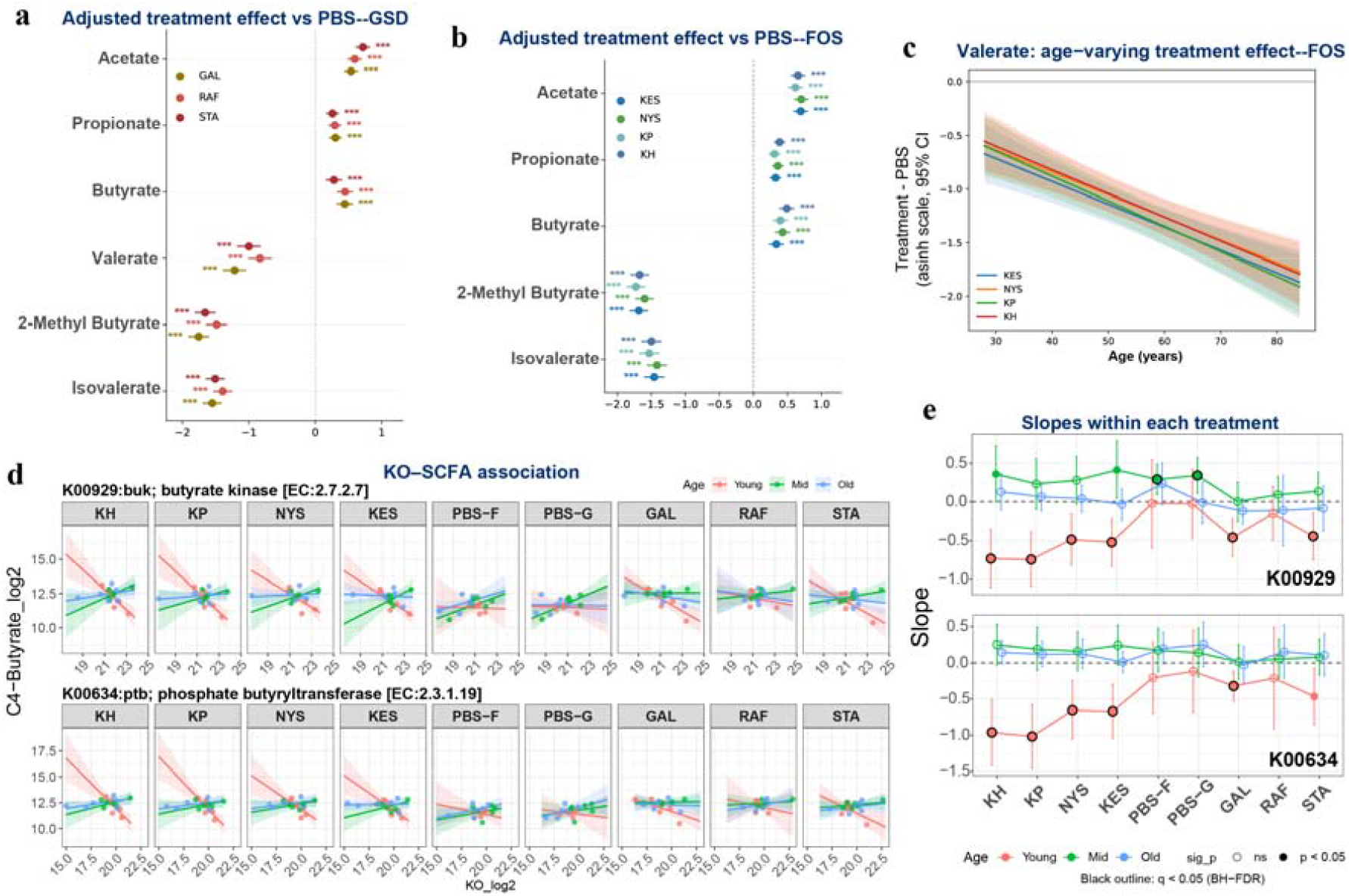
Age-dependent short-chain fatty acid (SCFA) responses to oligosaccharide treatments and KO–SCFA associations. (a, b) Age-adjusted treatment effects on selected SCFAs and BCFAs. Points show estimated treatment effects relative to PBS from the continuous-age mixed-effects model, and horizontal lines indicate 95% confidence intervals. Only metabolites with significant overall treatment effects and no treatment-by-age interaction are shown. Asterisks indicate adjusted significance levels (*\*\*\*q < 0*.*001*) (c) Age-varying treatment effects for C5-valerate, shown as estimated differences relative to PBS (Treatment − PBS) across age (years). Curves are model-predicted contrasts with 95% confidence bands. (d) KO–SCFA associations (LMM) between C4-butyrate (log_2_) and KO abundance (log_2_) within each treatment. Lines represent age-stratified fitted relationships (young/mid/old) with 95% confidence bands; points are subject-level observations. (e) Estimated KO–SCFA slopes within each treatment from the LMM, stratified by age group; significance codes indicate whether the slope differs from 0 after BH-FDR correction (*q < 0*.*05*).

Given the strong age-associated enrichment of *M. smithii*, a gut methanogenic archaeon, we next examined the corresponding methanogenesis-related functional signal. Under PBS conditions, we found a *M*.*smithii*-associated M00567 (KEGG module annotated as CO□-to-methane methanogenesis) signal was largely undetected in young and mid microbiomes but was consistently detected at higher abundance in old microbiomes in both oligosaccharide datasets (Fig. 6d). This pattern indicates that the elevated abundance of *M. smithii* in older microbiomes was accompanied by a detectable methanogenesis-related functional signature. We then examined whether oligosaccharide treatment altered this age-associated M00567 signal. Because M00567 was largely absent from young and mid microbiomes, treatment-associated changes were evaluated within the old group. In old microbiomes, both FOS and GSD treatments reduced M00567 abundance relative to PBS (Fig. 6e). Together with the taxonomic decrease in *M. smithii* after treatment, these results suggest that oligosaccharide exposure attenuated an *M. smithii* signature in microbiomes from older adults and its associated methanogenesis-related functional potential.

### Core SCFAs increase across treatments, while valerate and butyrate–pathway coupling show age- and context-dependent effects

We quantified lactate together with 10 fatty acid metabolites in the culture supernatant and found that acetate, propionate, and butyrate increased consistently across all oligosaccharide treatments relative to the control. In contrast, 2-methylbutyrate, valerate, and isovalerate showed a global decrease (Fig 7a). Notably, valerate exhibited a significant Treatment × Age interaction, while the main effect of Age was not significant after averaging across treatments; together, these results indicate that age primarily modulated the treatment response in a treatment-specific manner rather than imposing a uniform age trend across all conditions. Despite treatment-dependent effect sizes, we observed an overall tendency toward lower valerate levels with increasing age (Fig 7b, c).

Butyrate is an important microbial metabolite for host physiology and can be produced through terminal reactions involving butyrate kinase/phosphate butyryltransferase and CoA-transferase-associated enzymes. We therefore examined whether measured butyrate levels were associated with the abundance of enzymes annotated to these terminal steps. KO–SCFA association LMMs showed significant age-dependent abundance–butyrate associations for butyrate kinase (buk; K00929) and phosphate butyryltransferase (ptb; K00634) (Abundance × Age, FOS: buk: *p*≈*7*.*18 × 10*□ □; ptb: *p*≈*3*.*00 × 10*□ □; GSD: buk: *p*≈*5*.*68 × 10*□ □; ptb: *p*≈*1*.*73 × 10*□^*2*^). In contrast, the CoA-transferase-associated signal, assessed using K01034, showed weaker support, with significance observed only in the FOS set (FOS: *p*≈*2*.*72 × 10*□^*2*^; *GSD: p*≈*0*.*15*). Stratified slope analyses further showed that, in young subjects, both buk and ptb had negative abundance–butyrate slopes under multiple treatments (KH, KP, NYS, KES, GAL, and STA for buk only), with slopes significantly below zero after BH correction (BH-adjusted *q < 0*.*05*) (Fig. 7d, e). Residual diagnostics and LOSO sensitivity analyses supported the overall stability of the main buk- and ptb-associated slope patterns under oligosaccharide treatments, although a small number of subgroup-specific estimates showed limited single-donor sensitivity that did not alter the main interpretation (Supplemental Table 10 and 11; Supplemental Fig. 8 and 9). Pairwise age comparisons were consistent with this pattern (Supplemental Table 12). Although the K01034–butyrate slopes did not remain significantly different from zero after multiple-testing correction, the FOS treatments still suggested a positive trend (Supplemental Fig. 10). Collectively, these results suggest that the coupling between terminal butyrate-pathway enzyme abundance and extracellular butyrate varies with age and treatment.

## Discussion

Dietary non-digestible oligosaccharides are often treated as a single prebiotic class, but differences in their chemical structure can lead to distinct microbial responses and metabolic outcomes in the gut microbiome. In this study, we combined a controlled ex vivo fermentation system with deep DIA metaproteomics and targeted metabolomics across two glycan classes (FOS and GSD), seven related substrates, and three donor age groups. This design allowed us to separate conserved responses from donor-dependent contingencies and to ask not only whether oligosaccharides stimulate beneficial outputs, but how substrate structure and donor age shape upstream enzyme programs, functional allocation across producer guilds, and downstream metabolite readouts.

A central finding of this study is that oligosaccharide structure organizes a functional response landscape rather than simply separating substrates into broad prebiotic classes. Glycan structural distances were strongly associated with donor-level KEGG module response distances, indicating that structurally related glycans produced more similar functional responses overall. However, this relationship did not imply functional equivalence among closely related substrates. Within the FOS series, degree of polymerization was associated with 68 significant module-level response gradients, showing that chain-length differences within a single glycan family remained functionally resolvable. Thus, glycan structure shaped both the large-scale separation between glycan families and the finer graded responses among closely related FOS substrates.

This global structure–function relationship was reflected in primary carbohydrate-processing enzymes. Along the FOS chain-length series, both primary fructan-cleaving CAZymes increased with increasing degree of polymerization, consistent with a coherent structure–response relationship in which longer fructans sustain stronger or more persistent primary processing demands^30^. Because total supplied fructose residues increased only modestly under equal-mass addition, and the observed contrasts exceeded residue-only expectations, especially for fructosidase, this pattern supports a structure-dependent response beyond simple scaling by fructose input. In contrast, within the GSD set, linkage context and galactosyl decoration introduced a different form of structure dependence^31^. The RAF–STA difference in fructofuranosidase exceeded the expectation based on fructose-residue input alone. For alpha-galactosidase, the RAF–STA contrast did not clearly exceed the galactose-residue-based reference; however, because RAF and STA differ by only one galactosyl extension, this comparison cannot determine whether the higher signal under STA reflects residue load alone or a broader chain-length/structural-context effect. These results suggest that the community distinguishes substrates not only by the amount of relevant monosaccharide residues supplied, but also by glycosidic configuration, chain architecture, and residue context. Early carbohydrate-processing steps are therefore substrate-specific, and minor structural differences can change which enzymatic programs are prioritized.

The producer-level analyses further indicate that structure-responsive functions are organized through specific producer-guild contexts rather than representing only feature-level abundance changes. In the FOS series, the KO-level positive and negative DP-associated response sets defined by the structure–response model showed distinct producer behavior. The negative DP-associated KO set showed a stronger Bray–Curtis shift from matched PBS than expected from size-matched positive KO subsets, indicating that this producer compositional shift was not simply a consequence of the smaller negative KO set. This suggests that FOS chain length can organize not only the direction of functional responses, but also the producer contexts in which these responses occur.

This set-level result complements the targeted enzyme-level producer analysis. Two primary fructan-processing CAZymes were supported by distinct producer guilds, arguing against a single universal fructan degrader group. Fructosidase was dominated by a relatively stable set of *Bacteroidota*-associated producers, consistent with the well-established ability of *Bacteroides* to deploy dedicated carbohydrate utilization programs for complex glycans^32^. By contrast, β-fructofuranosidase showed more pronounced producer variability across glycan contexts and fewer active producers under RAF/STA compared with FOS. Ecologically, this pattern is compatible with the idea that FOS, containing multiple fructose units and chain-length variability, provide broader microbial access to fructose-linked intermediates^30,33^, whereas raffinose-family substrates contain a sucrose core with a single fructose and additional galactosyl decorations, which may narrow entry points for fructose-linked steps and constrain the active producer set^31^. Together, these observations support a niche-partitioning model where substrate structure determines which guild performs the initial deconstruction, while still allowing convergence on shared downstream fermentation outputs.

Beyond the FOS KO-set analysis, the RAF–STA comparison provided targeted mechanistic resolution for how a specific enzyme signal changed within the GSD series. The STA-associated decrease in beta-fructofuranosidase relative to RAF was larger than expected from fructose-residue input alone and was associated with a distinct WGCNA module context, motivating a producer-level decomposition of this signal. This decrease was not explained by reduced producer biomass or wholesale producer turnover, but instead by reduced functional output within largely shared producer sets. This attribution is important because it distinguishes two very different ecological interpretations: community restructuring versus reallocation of functional investment inside a stable producer cohort. In the RAF/STA context, the additional galactosyl residue in STA may shift functional demand toward alpha-galactose handling and associated uptake or catabolism, while reducing relative investment in fructose-linked cleavage steps.^31,34^. This type of decomposition, separating enzyme change into producer abundance, producer turnover, and per-producer output, offers a generalizable framework for interpreting inconsistent functional signals in multi-substrate microbiome studies.

At the network level, the primary-cleavage CAZymes were embedded within a broader, conserved physiological response. Although FOS and GSD yielded independent WGCNA networks, the target-containing modules converged on highly similar KEGG enrichment patterns, including positive enrichment for ribosome and amino-acid biosynthesis pathways and coordinated signals in secretion/export and chemotaxis-related functions. These patterns suggest that variation in first-cleavage capacity is coupled to a broader anabolic and growth-associated program^34,35^, rather than representing isolated enzymatic events. The metabolomics data reinforces this interpretation: multiple free amino acids increased across treatments, consistent with a community-level anabolic shift during oligosaccharide utilization. Conceptually, this supports a model in which prebiotic substrates trigger coordinated investment in biomass production, protein synthesis, and environment sensing/interaction, while downshifting selected catabolic or interconversion pathways when carbohydrate utilization dominates the community’s metabolic agenda^5,6^.

Within this broader program, we identified a robust cross-family association between primary fructan processing and tryptophan-associated microbial functions. Across both glycan families and both primary CAZymes, tryptophan synthase was consistently and positively associated in correlation space. This association was also reflected in donor-consistent differential expression patterns, with increased tryptophan synthase and decreased tryptophanase across most oligosaccharide treatments. In parallel, extracellular tryptophan levels were higher under oligosaccharide treatments than under PBS, indicating that oligosaccharide fermentation was accompanied by a coordinated shift in both tryptophan-associated enzyme profiles and the extracellular tryptophan pool. Producer profiling further showed that fructan-cleaving CAZymes and tryptophan synthase were assigned to largely distinct species-level producer groups, with only a small subset of species carrying both functions. This separation supports a guild-level organization in which primary fructan degradation and tryptophan biosynthesis-associated capacity are distributed across different members of the treatment-responsive community. Together, these patterns suggest that oligosaccharide fermentation reshaped microbial tryptophan-associated metabolism, involving increased biosynthesis-associated capacity, reduced tryptophan-degrading activity, and altered extracellular tryptophan availability. Although tryptophan synthase abundance and extracellular tryptophan were both higher under oligosaccharide treatments, the ex vivo system does not resolve the source of the extracellular tryptophan pool, particularly because tryptophan was already present in the culture medium. The observed increase could therefore reflect reduced microbial consumption or degradation, altered uptake or release, biomass-related changes, or new microbial synthesis. Nevertheless, changes in extracellular tryptophan availability are biologically relevant because tryptophan serves as a precursor for host metabolic and immune pathways, including serotonin and kynurenine metabolism^36,37^. Thus, these results suggest that oligosaccharide fermentation can reshape tryptophan-associated microbial functions and extracellular tryptophan availability, although whether these ex vivo changes influence host tryptophan metabolism or donor physiology will require in vivo validation with parallel host readouts.

Oligosaccharide treatments also appeared to shift community carbohydrate use away from host-derived glycans, as reflected by coordinated decreases in proteomic markers associated with mucin glycan degradation and lower mucin-chain scores. The magnitude of this shift differed across treatments, with GAL, KES, and NYS showing the strongest downshifts, RAF and STA the weakest, and KP and KH occupying intermediate positions. Within the FOS series, responses tended to be stronger for the shorter-chain compounds KES and NYS than for the longer-chain compounds KP and KH, although these within-family differences were not statistically significant. More importantly, comparisons among the short-chain treatments showed that KES, NYS, and GAL all produced stronger downshifts than the RFO compounds RAF and STA, indicating that the observed differences cannot be explained by chain length alone. These results suggest that both degree of polymerization and other structural features, such as family type and glycosidic configuration, may influence how exogenous oligosaccharides alter microbial investment in mucin glycan degradation. In a host context, reduced investment in mucin glycan degradation could be compatible with lower pressure on the mucus layer, consistent with prior work linking fiber deprivation to increased mucus degradation^38^. However, such implications remain indirect in the present ex vivo setting, and confirmation will require direct assessment of mucus integrity, epithelial responses, and inflammatory outcomes.

Age emerged as a modifier of oligosaccharide responsiveness, influencing the baseline taxonomic and functional context in which fermentation responses occurred rather than simply changing the direction of core metabolite outputs. A prominent example was the enrichment of *M. smithii*, a prevalent non-pathogenic methanogenic archaeon in the human gut, in the microbiomes of older adults. This finding is consistent with recent shotgun metagenomic evidence showing age-associated dynamics of gut methanogenic archaea and an increased prevalence of high-methanogen phenotypes defined by *Methanobrevibacter* abundance with age^39^. *M. smithii* obtains energy through methanogenesis and, in culture, primarily reduces CO□ to methane using H□, with formate used to a lesser extent^40^. This biology provides a rationale for examining M00567, a KEGG module annotated as CO□-to-methane methanogenesis. In our dataset, the higher abundance of *M. smithii* in older donors was accompanied by higher detection and abundance of M00567, whereas this signal was largely absent from young and mid-life donors. Oligosaccharide treatment reduced both *M. smithii* abundance and the M00567 signal in older microbiomes, suggesting attenuation of an age-associated *M. smithii*-linked methanogenesis-related signature. Since methane production was not directly measured, we interpret this as a methanogenesis-related taxonomic and functional signature rather than direct evidence of altered methane output. Because *M. smithii* is a common gut archaeon with normal ecological roles as well as context dependent clinical associations, including enrichment in constipation associated settings and depletion in ulcerative colitis^40-42^, these results do not support a simple beneficial-versus-harmful interpretation. Instead, they indicate that oligosaccharide exposure reduced an elevated *M. smithii* associated functional signature in older microbiomes.

Finally, across all oligosaccharide treatments we observed a consistent rise in the major fermentation SCFAs (acetate, propionate, butyrate) together with decreases in valerate and several BCFAs often associated with proteolytic fermentation^43,44^. This pattern is broadly consistent with the canonical prebiotic model: fermentable carbohydrates promote saccharolytic fermentation and shift community metabolism toward main SCFA production, a central route by which diet–microbiome interactions can influence donor physiology^45,46^. FOS induced a progressively stronger reduction in valerate relative to PBS with increasing donor age. Because valerate has been reported to support epithelial barrier properties in vitro^47^, this shift may have potential relevance to barrier-related metabolism. As well, valerate can also arise from protein- or amino acid-linked fermentation and may have context-dependent effects in aging-associated inflammatory settings^48^. Together, these results suggest that FOS treatment is associated with an age-dependent reduction in valerate, pointing to differential remodeling of minor SCFA metabolism in older microbiomes. Beyond this change in minor SCFA metabolism, the butyrate analysis showed that oligosaccharide-associated butyrate accumulation was accompanied by age specific enzyme–metabolite relationships. Although butyrate increased consistently across treatments, the relationship between terminal butyrate-pathway enzyme abundance and extracellular butyrate was not uniform across donor age groups. In particular, buk and ptb showed the strongest age-dependent abundance–butyrate associations, with negative slopes in young donors under several oligosaccharide treatments. This indicates that, despite the overall increase in butyrate after oligosaccharide treatment, variation in buk/ptb abundance did not explain the magnitude of extracellular butyrate accumulation in a uniform manner. This pattern suggests that total butyrate output and pathway-associated enzyme abundance capture related but distinct aspects of the fermentation response. Extracellular butyrate represents a net metabolic outcome, whereas enzyme abundance also reflects the producer taxa, their growth state, substrate use, and potential downstream consumption or cross-feeding. The CoA-transferase-associated signal assessed using K01034 was weaker and less consistent, showing only a positive tendency under FOS treatments. Together, these findings indicate that similar increases in butyrate concentration can be accompanied by different underlying enzyme–metabolite relationships across age groups, supporting an age-aware interpretation of fermentation outputs.

This study has limitations that define clear next steps. First, the ex vivo fermentation system isolates microbial metabolism from host absorption, immune signaling, and intestinal physiology. Therefore, host-facing interpretations of the observed *M. smithii*-associated M00567 signature, extracellular tryptophan availability, valerate remodeling, and SCFA responses will require in vivo validation with direct physiological readouts, such as breath methane, bowel-function endpoints, and epithelial or immune markers. Second, the relatively small number of donors in each age group limited statistical power to resolve smaller or more variable effects, even though the study was sufficient to detect robust differences associated with substrate structure and donor age. Third, the short culture duration captures acute responses rather than long-term adaptation. Future studies incorporating time-series sampling, gas measurements, and isotope- or flux-based approaches would help distinguish pathway activity, substrate use, metabolite turnover, and microbial growth effects. Despite these constraints, by integrating structurally resolved oligosaccharides, donor age stratification, high-resolution metaproteomics, and metabolite readouts, our work provides a framework in which oligosaccharide architecture shapes upstream primary-cleavage guilds and coordinated functional responses, while donor age influences the baseline taxonomic and functional context in which these responses emerge.

## Conclusion

This study provides a structure- and donor-aware framework for interpreting prebiotic responses in the gut microbiome. Using matched ex vivo cultures, DIA metaproteomics, and targeted metabolite profiling across age groups, we show that oligosaccharides structural distance organizes donor-level functional response divergence, while closely related oligosaccharides remain functionally distinguishable rather than interchangeable. Within the FOS series, chain length was associated with graded module- and KO-level response patterns and progressively stronger induction of primary fructan-cleaving CAZymes. Producer-level analyses further showed that FOS DP-responsive KO sets and GSD-associated enzyme changes were linked to distinct producer contexts, including producer compositional shifts for negative FOS DP-associated KOs and altered functional investment within shared producer backgrounds for the RAF–STA beta-fructofuranosidase contrast.

Beyond carbohydrate cleavage, oligosaccharide treatments were associated with a conserved anabolic functional program, altered tryptophan-associated microbial functions, higher extracellular tryptophan availability, and reduced mucin glycan degradation-associated markers. These findings indicate that oligosaccharide responses involve broader shifts in microbial resource allocation, amino acid associated metabolism, and host-glycan foraging potential. Donor age did not uniformly change the direction of core fermentation outputs, but it modified selected taxonomic and functional axes, including an age-associated *M. smithii* and M00567 methanogenesis-related signature, valerate responses, and butyrate pathway–metabolite coupling. These results argue that precision prebiotic strategies should not be evaluated only by total metabolite changes, but also by substrate-specific enzyme programs, the producer guilds carrying them, and donor-dependent links between microbial function and metabolic output.

Because this work was performed ex vivo, future studies should validate these structure- and age-linked patterns in vivo. Together, our findings show that oligosaccharide architecture shapes upstream primary-cleavage guilds and coordinated functional responses, while donor age influences the baseline taxonomic and functional context in which these responses emerge.

## Data availability

The mass spectrometry proteomics data have been deposited to the ProteomeXchange Consortium *via* the PRIDE^49^. Partner repository with the dataset identifier: PXD079344.

## Author contributions

A.Z. designed and performed most experiments, processed and analyzed the data, prepared the figures, and wrote the original draft. Q.W. processed the data and contributed to data analysis, figure visualization, and metabolite sample preparation. H.Q. performed the amino acid and SCFA assay procedures. J.M. designed and supervised the study, contributed to conceptualization and methodology, and reviewed and revised the manuscript. Z.N. was responsible for mass spectrometry operation. C.E.R. performed part of the amino acid assay procedures. D.F. designed and supervised the project, contributed to conceptual development and methodology, secured funding, and reviewed and revised the manuscript.

## Acknowledgements

This study received funding from the NSERC-CREATE Technologies for Microbiome Science and Engineering (TECHNOMISE) program under grant number CREATE-497995-2017, which was awarded to D.F. A.Z. was supported by a scholarship from the China Scholarship Council (CSC). A.Z. also acknowledges financial support from Quadram Institute Bioscience for a research visit related to this study. The authors gratefully acknowledge the assistance of ChatGPT, Grammarly, DeepL and Gemini in enhancing the English language presentation of this manuscript. The instrument illustration in workflow was generated with the assistance of ChatGPT and Gemini.

